# Hybrid deep learning–mechanistic modeling of cellular dynamics from a spatiotemporal single-cell atlas

**DOI:** 10.1101/2025.10.02.679951

**Authors:** Xiaoyu Duan, Vipul Periwal

## Abstract

Single-cell measurement technologies provide a powerful framework for studying cellular heterogeneity, transitions, and regulatory networks, yet reconstructing the underlying dynamical processes governing these transitions remains a major challenge due to the high dimensionality of gene expression data. To address this, we develop a variational autoencoder (VAE)–latent neural ordinary differential equation (ODE) approach that learns a low-dimensional latent representation of cellular states and models their temporal evolution. We apply our framework to the single-cell fluorescence imaging spatiotemporal atlas of Drosophila melanogaster blastoderm embryos via spatial registration, which comprises six registered developmental time points prior to gastrulation. In our approach, gene expression profiles are encoded into a latent space where a neural ordinary differential equation (ODE) is trained to capture the continuous dynamics of cellular states over time, and a decoder subsequently maps these evolving latent representations back to the original high-dimensional gene expression space, enabling accurate reconstruction of observed transcriptional patterns. While such black-box deep learning approaches excel at capturing complex dynamical trajectories, they are inherently limited in their ability to predict the effects of combinatorial perturbations. To overcome this limitation, we leverage the inferred latent dynamics as a foundation for fitting a mechanistic Hill-function model of gene regulation, which—guided by the black-box representations—enables interpretable predictions by systematically perturbing couplings between specific gene pairs and thus provides mechanistic insight into developmental regulatory programs.

**Author summary:** We set out to learn how genes interact over time during early development using data measured from individual cells. Many powerful deep learning models can fit complex patterns, but they are hard to edit or use for “what-if” genetic changes. Classical mechanistic models are easier to interpret but struggle when the data are sparse. We built a hybrid approach that combines both strengths.

First, we trained a deep learning model to place each cell’s gene activity into a smooth, low-dimensional space and to estimate how these states change over time. We then used those estimates to fit a simple, editable model of gene regulation based on standard biochemical response curves. This second model lets us turn specific regulatory connections on or off and predict the effects without retraining the whole system.

Using a spatiotemporal atlas of the fruit fly embryo, we show that the hybrid model makes accurate short-term predictions and yields clear, testable hypotheses about which genes control others. Our study demonstrates a practical way to turn flexible deep learning into interpretable, mechanistic insight for dynamic single-cell data.

## Introduction

One key challenge in systems biology is to learn interpretable, continuous-time dynamics from high-dimensional single-cell snapshots. Another challenge is to endow such models with mechanistic structure so they can generalize to combinatorial perturbations and support causal reasoning. Addressing the first challenge, three major families of dynamic methods have emerged in single-cell analysis: (i) *Latent ordinary differential equations (ODEs)* that couple deep generative encoders/decoders with continuous-time latent dynamics (1; 2); (ii) *Dynamic optimal transport/continuous flows* that learn population-level trajectories between time-stamped snapshots (e.g., TrajectoryNet) (3; 4); and (iii) *Velocity-based methods* that infer gene- or cell-wise velocities from spliced/unspliced reads and reconstruct transcriptomic vector fields with fate mapping (scVelo, CellRank, dynamo) (5; 6; 7; 8). In parallel, ODE-based GRN inference (e.g., SCODE, GRISLI) imposes sparsity directly in gene space (9; 10).

Meeting the second challenge requires moving beyond black-box accuracy. Deep models can fit complex trajectories, yet they extrapolate poorly to unseen *combinations* of interventions: with 10 genes and three states per gene (activation, inhibition, no change), the design space is 3^10^, making exhaustive black-box simulation impractical. At the same time, mechanistic ODEs can support structured what-if analyses but typically require reliable temporal derivatives, which are hard to estimate from coarsely sampled data. We therefore propose a hybrid approach: first train a VAE with a latent neural ODE to interpolate sparse time points and produce variance-reduced gene-space velocities via the decoder pushforward; then fit a sparse, editable Hill-function ODE to these velocities and short-horizon predictions, combining deep learning’s accuracy with the interpretability and combinatorial generalization of an explicit dynamical system.

We demonstrate our approach on a quantitative single-cell fluorescence imaging atlas of Drosophila melanogaster embryos (11), not on single-cell RNA sequencing; hence no splicing kinetics or count-based RNA velocities are used. This dataset, registered into a common coordinate system, forms a spatiotemporal atlas of early embryonic development and provides cellular-resolution measurements of gene expression across multiple time points. It has been used in prior studies, including mechanistic modeling efforts such as (12), making it a compelling test case. However, the dataset’s limited temporal resolution and missing gene coverage pose challenges for directly learning continuous gene dynamics, motivating our two-stage modeling approach. Full details are provided in the Data section.

(12) applied an outlier-insensitive regression approach—erf-weighted least absolute deviation (LAD)—to this same dataset to directly infer a mechanistic gene network in the 99-dimensional expression space without using spatial information. Their model incorporated both linear and quadratic terms, enabling one-step-ahead predictions of discrete gene dynamics and allowing simulation of knockout and perturbation scenarios, which showed qualitative agreement with experimental results. While their approach demonstrated robustness to outliers and competitive predictive accuracy relative to black-box neural networks, it relied on discrete-time regression rather than continuous latent-space dynamics. In this work, we extend this line of investigation by introducing a hybrid VAE–neural ODE–Hill function pipeline that uses latent continuous dynamics to generate smooth derivative estimates for mechanistic model fitting.

Related dynamic-inference methods in single-cell analysis include RNA velocity and its dynamical extensions (5; 6; 8), and optimal-transport–based trajectory reconstruction (4). Our approach differs in using a learned continuous latent vector field as a teacher for fitting a sparse, editable mechanistic ODE in gene space. Deep generative models such as single-cell variational inference (scVI) (13) and continuous-time neural model (2) motivate our “VAE with latent neural ODE” design.

To model gene dynamics, we adopt a mechanistic Hill-type ODE in which each gene’s production is a saturating, monotone function of its regulators, opposed by linear decay. The interaction matrix encodes directed gene–gene influences; we impose an *ℓ*_1_ penalty (Least Absolute Shrinkage and Selection Operator, Lasso) during training to induce sparsity, reflecting that real regulatory networks are not fully coupled. We use Hill nonlinearities because they yield bounded, well-behaved vector fields that integrate stably over the horizons we study. By contrast, more flexible surrogate representations (e.g., symbolic regression without explicit stability constraints) can produce stiff or even divergent trajectories when extrapolated, complicating long-horizon integration.

We therefore adopt a two-stage strategy. First, we train a variational autoencoder (VAE) to compress the original 99-dimensional spatiotemporal gene-expression data into a low-dimensional latent state. We then learn a latent-space neural ODE that captures smooth continuous dynamics over this latent state. We push the latent vector field through the decoder, yielding an induced ODE for the genes that we integrate directly while holding spatial coordinates fixed. These decoder push-forward velocities provide locally smooth targets and short-horizon predictions used to fit the explicit, sparse Hill-type model, leveraging the expressivity of neural ODEs for interpolation and the interpretability of a mechanistic ODE for downstream analysis.

To orient the reader, the Data section describes the atlas and preprocessing. The VAE section details the variational autoencoder and mask-respecting training, and the Imputation section presents the fixed-point procedure. The Neural ODE section introduces the latent Neural ODE and the decoder push-forward construction of gene-space dynamics, and the Hill-model section specifies the sparse Hill-network model and its estimation. The Results then compare teacher-guided versus data-only Hill fitting, examine spatial universality under region-restricted supervision, and demonstrate interpretable perturbations using the trained VAE, neural ODE, and the Hill model—both by mathematically knocking out specific genes and by decoupling selected gene pairs in the Hill model. We conclude with a Discussion of limitations, implications, and future directions.

## Materials and methods

### Data

To run this process, we use the dataset from (11) as an example. This dataset was constructed by applying a registration technique to align data from 1822 *Drosophila melanogaster* blastoderm embryos into a unified spatiotemporal atlas across 6,078 embryo positional bins. It includes expression measurements for 95 mRNAs and 4 proteins, collectively referred to as 99 “genes” for brevity throughout this paper. These measurements span the 50-minute period preceding gastrulation, uniformly divided into six time points, labeled as *t* = 0 to *t* = 5. Out of the 99 genes, 27 of them have complete measurements across all six time points, while the remaining 72 are primarily measured during the last three time points closer to gastrulation. In the following sections, we detail the application of our framework to impute the missing entries in this dataset. Following (12), we use a 90/10 positional-bin split with within-split shuffling, holding out a fixed test set of 608 bins randomly sampled across the embryo for all evaluations unless otherwise noted.

### Methods at a glance

We use an eight-step pipeline that couples a learned latent “teacher” with a sparse mechanistic “student” (Fig 1).

**Fig 1.**
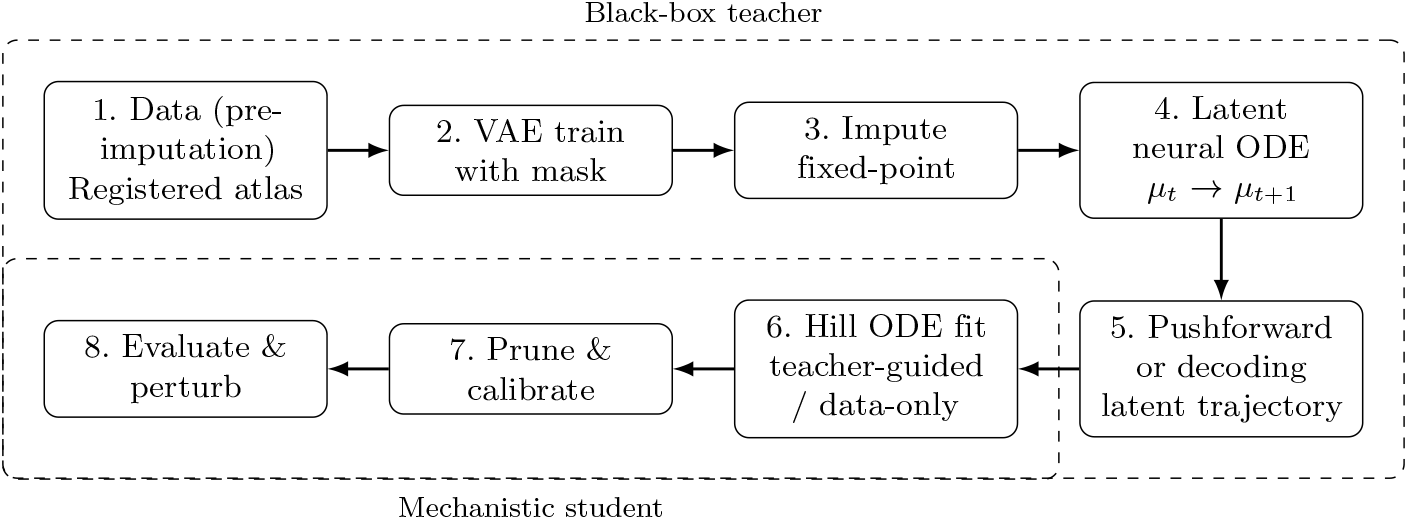
Eight-step pipeline: (1) data (pre-imputation); (2) mask-respecting VAE; (3) fixed-point imputation; (4) latent neural ODE; (5) compute teacher derivatives; (6) sparse Hill ODE fit (teacher-guided or data-only); (7) pruning and calibration; (8) evaluation and perturbations.

1. Data (pre-imputation): load the registered blastoderm atlas and apply an initialization scheme for all missing gene entries so that the encoder always consumes complete vectors. These pre-imputed values serve only as placeholders for valid inputs and are subsequently refined by the fixed-point imputation.
2. VAE training with a mask: train the autoencoder on genes plus spatial coordinates, but ensure that unobserved entries are masked out so they do not contribute to the reconstruction loss or parameter updates; only measured values influence learning.
3. Fixed-point imputation: iteratively fill missing gene entries with VAE predictions, keeping measured entries unchanged, until convergence.
4. Latent neural ODE: learn smooth continuous dynamics between successive latent states using an adaptive ODE solver.
5. Teacher derivatives in gene space: either by decoder pushforward, which computes gene-space velocities by multiplying the decoder Jacobian with the latent ODE vector field, or decode a dense latent ODE trajectory and compute finite-difference slopes on that path.
6. Hill ODE fitting: train a sparse gene-regulatory model using teacher-guided derivatives or, in a data-only regime, finite differences from observed time points.
7. Prune and calibrate: sparsify the interaction matrix, fix the surviving structure, and recalibrate remaining parameters for stable rollouts.
8. Evaluate and perturb: assess prediction error on held-out bins and across genes, test spatial generalization, and run targeted knockouts or edge edits to probe causal pathways.

### Variational Autoencoder (VAE)

Direct training of neural differential equation models in the full 99-dimensional gene expression space is computationally challenging. To address this, we trained a variational autoencoder (VAE) (14) to learn a compact low-dimensional latent representation of the data. Because the neural ODE operates in this latent space, we scaled the posterior standard deviation by a small constant factor to reduce overlap between the latent distributions of distinct samples.

Temporal information was not explicitly provided to the VAE during training; sample order was randomized, although the original time indices were retained for downstream analyses. In contrast, spatial context proved essential: excluding spatial coordinates during training led to latent representations poorly suited for dynamical modeling. Accordingly, the VAE input comprised both the 99 gene expression values and the three-dimensional spatial coordinates of the corresponding positional bins, while the decoder was trained to reconstruct only the gene expression values, not the spatial coordinates.

A binary mask identified observed genes per sample so that reconstruction was computed only over observed entries during training. The VAE loss is the sum of five components:

1. Masked reconstruction: mean-squared error over observed genes only.
2. KL divergence with warm-up: regularizes the posterior toward a standard normal and increases its weight early in training.
3. Smooth sparsity: gentle penalty on linear-layer weights that encourages sparser, simpler parameterizations.
4. Decoder smoothness: Jacobian-based proxy that discourages abrupt changes in the latent-to-gene mapping and is ramped up late in training.
5. Directional alignment: aligns local decoder changes with empirical temporal differences when a valid next time point is available.

The last three terms can slightly increase raw reconstruction error, but this bias toward simpler and smoother mappings is deliberate. An overfit black box with sharp, sample-specific fluctuations between adjacent time points provides a poor teaching signal for the downstream mechanistic model; its derivatives and trajectories are noisy, making the student harder to train and, in some cases, yielding worse fits than training the mechanistic model without a black-box teacher. To manage this trade-off, we prioritize checkpoints that balance accuracy and smoothness using a smoothness-aware validation criterion, which improves learnability for the mechanistic stage. Detailed definitions of the loss terms and training settings are provided in the Supplementary S1.

To visualize the structure of the latent space, we fit a Uniform Manifold Approximation and Projection (UMAP) embedding on the latent encodings of all positional bins and then fixed this embedding across time. Fig 2 shows the result: the left panel depicts the blastoderm embryo layout, and the right panel shows the same latent embedding colored by time point (*t* = 0 to *t* = 5). This representation highlights how cells at distinct developmental stages are separated yet connected within a continuous latent manifold, illustrating that the VAE has captured both spatial and temporal structure in a form suitable for downstream dynamical modeling.

**Fig 2.**
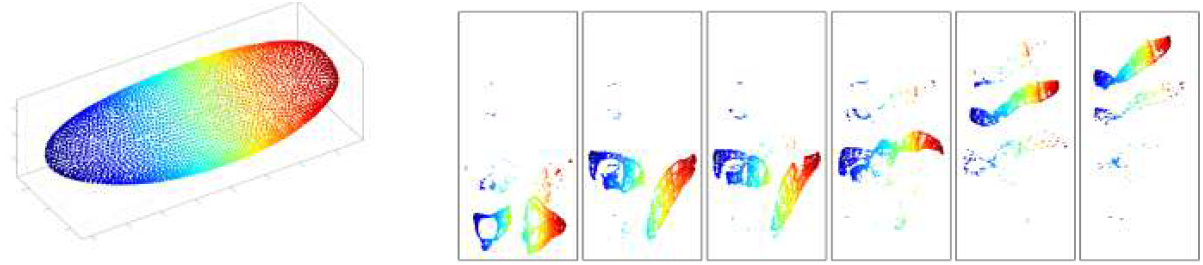
Embryo layout and UMAP embedding of latent states. Left: schematic of the blastoderm embryo with 6,078 spatial bins. Right: UMAP visualization of latent encodings across all bins, shown separately for the six developmental time points (*t* = 0 to *t* = 5, left to right). Red corresponds to posterior bins of the virtual embryo and blue corresponds to anterior bins.

### VAE-based imputation of missing gene expression

We used the VAE both to compress the full spatiotemporal gene profiles into a low-dimensional latent space and to impute unmeasured gene values. Because the encoder always consumes the complete gene vector, missing entries are supplied with placeholders to form valid inputs; during VAE training these placeholders are masked and do not contribute to the reconstruction term. Imputation is performed only after training: we apply the trained VAE in a fixed-point loop that repeatedly encodes the current gene vector together with spatial coordinates, decodes to predictions, and replaces only masked entries—leaving measured values unchanged—until changes at the masked positions are negligible or a preset iteration cap is reached. We experiment over three initialization schemes for missing entries.

1. Zero-fill: set missing gene expression entries to 0 at the current time.
2. Mean-fill: for each positional bin and gene, set missing gene expression entries in that bin to the mean of that gene’s observed values in that bin (per-gene, per-cell mean).
3. Random-fill: initialize each missing entry by a uniform draw within the observed range of that gene.

Fig 3 summarizes test-set reconstruction performance for the 99-gene decoder as a function of latent dimensionality. We compare three imputation initialization schemes (zero, mean, random) under two training objectives: the full five-term loss and a reduced objective that omits the decoder-smoothness and directional alignment components. In both settings, random-fill initialization consistently achieves the lowest mean absolute error (MAE), mean squared error (MSE), and the highest mean coefficient of determination (*R*^2^), while zero-fill initialization performs worst across dimensions. Unless otherwise noted, we therefore adopt the random-fill initialization in all subsequent experiments across latent dimensionalities.

**Fig 3.**
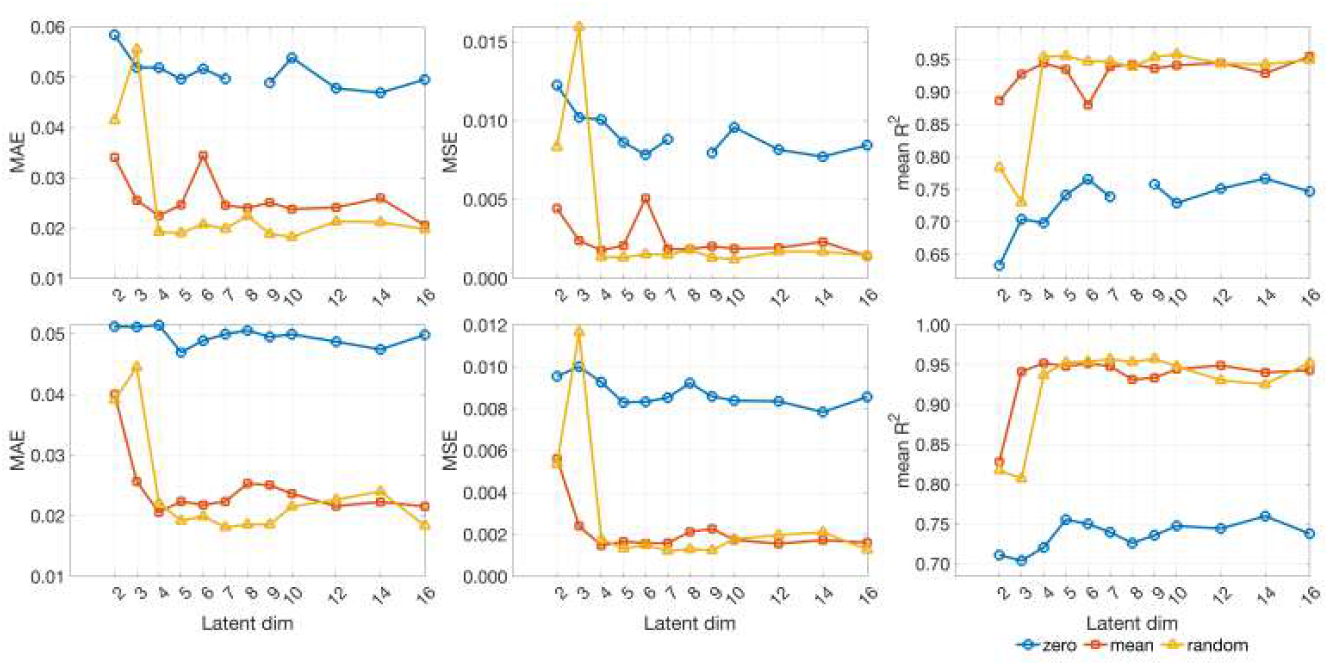
VAE reconstruction accuracy versus latent dimension on the testing set. Rows correspond to training objectives: top row uses the full five-term loss (reconstruction, KL, sparsity, decoder smoothness, directional alignment); bottom row removes the decoder smoothness and directional alignment components, retaining only reconstruction, KL divergence, and smooth sparsity. Columns show mean absolute error (MAE), mean squared error (MSE), and mean coefficient of determination value (*R*^2^). Curves compare imputation initializations (zero/mean/random). Across latent dimensionalities, random-fill consistently attains the lowest error and highest *R*^2^, while zero-fill is worst, under both training objectives. The zero-initialization point at latent dimension 8 is an outlier and is omitted from the plots.

### Neural ODE for Latent and Gene-Space Dynamics

We model state evolution in the VAE latent space with a continuous-time latent ODE. Let the latent state be 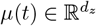, where *d*_*z*_ is the latent dimensionality. The dynamics are

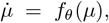

with *f*_*θ*_ a fully connected neural network; see (2) for neural ODE foundations. Training tuples (*µ*_*t*_, *µ*_*t*+1_, *σ*_*t*+1_) are constructed from VAE encodings of the imputed tables: *µ*_*t*_ and *µ*_*t*+1_ are latent means at successive time points for the same positional bin, and *σ*_*t*+1_ is the per–latent-dimension posterior standard deviation at the next time point.

For each tuple we integrate the ODE over a single interval starting from *µ*_*t*_ to obtain an endpoint 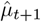, and learn parameters by minimizing a variance-normalized terminal error

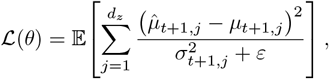

which places greater weight on well-determined latent directions; *ε >* 0 ensures numerical stability. Each interval is fit independently to avoid compounding integration error during training. We use an adaptive Dormand–Prince solver (RK45) for forward integration and the adjoint method for backpropagation, which controls memory usage with respect to the number of solver steps (15).

Although downstream analyses operate in gene space, the neural ODE is trained entirely in latent space. This decoupling avoids the instability and added complexity of end-to-end training of the VAE and dynamics, yet yields a smooth latent vector field that captures temporal structure.

#### Two black-box realizations used throughout

Let *g* ∈ ℝ ^99^ denote the gene vector and *s* ∈ ℝ^3^ the fixed spatial coordinates for a positional bin. The encoder consumes the concatenated input [*g* ∥ *s*] to produce *µ* = Enc([*g* ∥ *s*]), and the decoder maps latent states to genes, Dec: 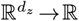 ^99^. We employ two ways to realize gene-space dynamics from the learned latent field:

1. **Pushforward gene-space ODE (“pushforward velocities”)**. The latent vector field is pushed through the decoder Jacobian to induce a gene-space vector field,

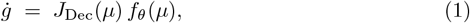

where *J*_Dec_(*µ*) is the decoder Jacobian with respect to the latent variables. Spatial coordinates *s* are held fixed and are not integrated. We integrate this gene-space ODE with the same solver setup described above. Because integration segments begin from the observed *g*(*t*_*k*_), this construction eliminates reconstruction error at each segment start and aligns the learned dynamics directly with gene-level readouts.
2. **Decoded latent rollouts**. Alternatively, we integrate the latent ODE to obtain *µ*(*t*) and decode the entire trajectory,

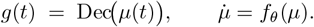 This realizes trajectories in gene space by decoding solutions of the latent dynamics. Unlike the pushforward formulation, decoded rollouts can inherit reconstruction error at segment endpoints.

These two realizations (pushforward gene-space ODE vs. decoded latent rollouts) are what we refer to as the *black-box dynamics* throughout the paper. This construction does not yield a pure vector field defined everywhere in gene space: because the encoder consumes both gene expression values and the fixed spatial coordinates of each bin, the induced derivatives are tied to specific spatial contexts rather than to arbitrary points. In practice, the black-box teacher provides reliable velocities only at the atlas bins, but these locally consistent derivatives still supply a valuable training signal for fitting the mechanistic Hill model compared with relying solely on finite-difference slopes.

### Mechanistic Hill-network model

Hill-type ODEs have also been successfully applied in classical studies of *Drosophila* gap-gene dynamics, such as the dynamic control framework of (16), motivating our use of this formulation here. We represent gene dynamics with a Hill-type regulatory ODE over *G* genes. Let 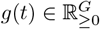 be the gene-expression vector. For each gene *i*,

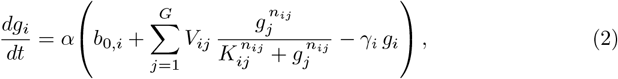

where *α >* 0 is a global production scale, *b*_0,*i*_ are basals, *V*_*ij*_ are directed interaction weights (activation if *V*_*ij*_ > 0, repression if *V*_*ij*_ < 0), (*K*_*ij*_, *n*_*ij*_) are Hill midpoints and exponents, and *γ*_*i*_ > 0 are linear decays.

To promote interpretability, we impose an *ℓ*_1_ penalty on *V* and hard-prune small entries after fitting. The surviving interaction support

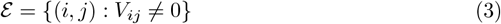

defines a directed gene–gene graph. In the Results calibrations, we keep both *ε* and *V* fixed and adjust only *α, γ, b*_0_, and (*K*_*ij*_, *n*_*ij*_) on (*i, j*) ∈*ε*.

We fit this Hill field either with or without guidance from a learned continuous model. The learned model, referred to as the black-box teacher, consists of a VAE encoder/decoder paired with a latent neural ODE. Consistent with the Neural ODE section, we form teacher gene-space derivatives in two ways: (i) pushforward gene-space ODE, where the latent vector field is mapped through the decoder Jacobian to obtain a velocity at the observed states (1) and we integrate piecewise starting from the measured *g*(*t*_*k*_); and (ii) decoded latent rollouts, where we integrate the latent ODE across each interval, decode a dense path in gene space between *t*_*k*_ and *t*_*k*+1_, and compute derivatives along that path by finite differences on a fine grid. In both constructions the spatial coordinates *s* are held fixed and are not integrated. When either of these teacher derivatives is used during training we call the regime teacher-guided; when neither is used and derivatives come only from observed finite differences at the original sampling times, we call the regime data-only. These names are used in the Results section to compare performance under matched conditions.

Estimation uses a simple neural-network training objective that minimizes a weighted sum of two terms: a derivative-matching loss that aligns the Hill right-hand side with teacher-informed velocities (teacher-guided) or finite-difference estimates (data-only), and a one-step prediction loss obtained by integrating the ODE from *t* to *t* + 1 (window *W* = 1). The weights are scheduled from derivative-heavy early epochs to prediction-heavy late epochs. Details are provided in the Supplementary S1.

## Results

### Impact of teacher guidance on mechanistic Hill-model fitting

We next ask whether using the learned “black-box teacher” introduced in the Neural ODE section improves downstream mechanistic fitting relative to relying on observed data alone. Because the teacher is optimized for its own reconstruction and integration objectives, there is no guarantee that it provides a superior signal for training a Hill-network ODE. We therefore compare, under matched conditions, the *teacher-guided* regime (VAE with a latent neural ODE providing gene-space velocities) against the *data-only* regime (no guidance), evaluating identical one-step forecasts (*t* → *t*+1) on a fixed test set.

We restrict to *G*=27 genes with complete measurements across all six time points to avoid imputed ground truth and to keep supervision windows identical across regimes. Both pipelines enforce the same sparsity and interpretability constraints: an *ℓ*_1_ penalty on the interaction matrix *V*, hard pruning of small entries, and a fixed interaction structure thereafter. After pruning, we freeze the interaction support (3) (the directed gene–gene graph) and the coefficients *V*_*ij*_; a subsequent calibration stage then tunes only *α, γ, b*_0_, and the Hill shape parameters (*K*_*ij*_, *n*_*ij*_) for (*i, j*) ∈ *ε* without altering *ε* or *V*. Predictions are obtained by integrating the ODE over a single interval.

Results for *G*=27 genes (window *W* =1) show that *teacher-guided* fits achieve equal or lower error after the identical calibration sequence, with the largest advantage under stronger sparsity. Final errors and surviving edge counts are summarized in Table 1.

**Table 1.** One-step forecast errors and nonzeros after pruning (|*V*_*ij*_| < 10^*−*3^). “Non-zero edges” counts |*ε* | = # *{* (*i, j*): *V*_*ij*_ ≠ 0 *}*. Teacher-guided models are as good or better; the advantage widens under stronger sparsity.

| Setting | Regime | MAE with SD | MSE | Non-zero edges |
| --- | --- | --- | --- | --- |
| $\lambda_{\ell_1}=10^{-3}$ | Teacher-guided | **0.05716**$\pm 0.02134$ | **0.00716** | 151 |
| $\lambda_{\ell_1}=10^{-3}$ | Data-only | 0.05958$\pm 0.02209$ | 0.00775 | 15 |
| $\lambda_{\ell_1}=10^{-4}$ | Teacher-guided | **0.05239**$\pm 0.01820$ | **0.00592** | $\approx 232$ |
| $\lambda_{\ell_1}=10^{-4}$ | Data-only | 0.05279$\pm 0.01893$ | 0.00600 | $\approx 289$ |

Following the definitions in Neural ODE section, Fig 4 contrasts the two black-box realizations over each interval [*t*_*k*_, *t*_*k*+1_]: (A) the *pushforward gene-space ODE* (1) *integrated from the observed g*(*t*_*k*_) (blue), and (B) the *decoded latent trajectory* obtained by decoding a path in latent space (red, dashed). For Fig 4 only, we ablate the latent dynamics in (B) and use linear interpolation between *µ*(*t*_*k*_) and *µ*(*t*_*k*+1_) before decoding. This shows how the VAE itself performs where we set latent dynamics to be trivial.

**Fig 4.**
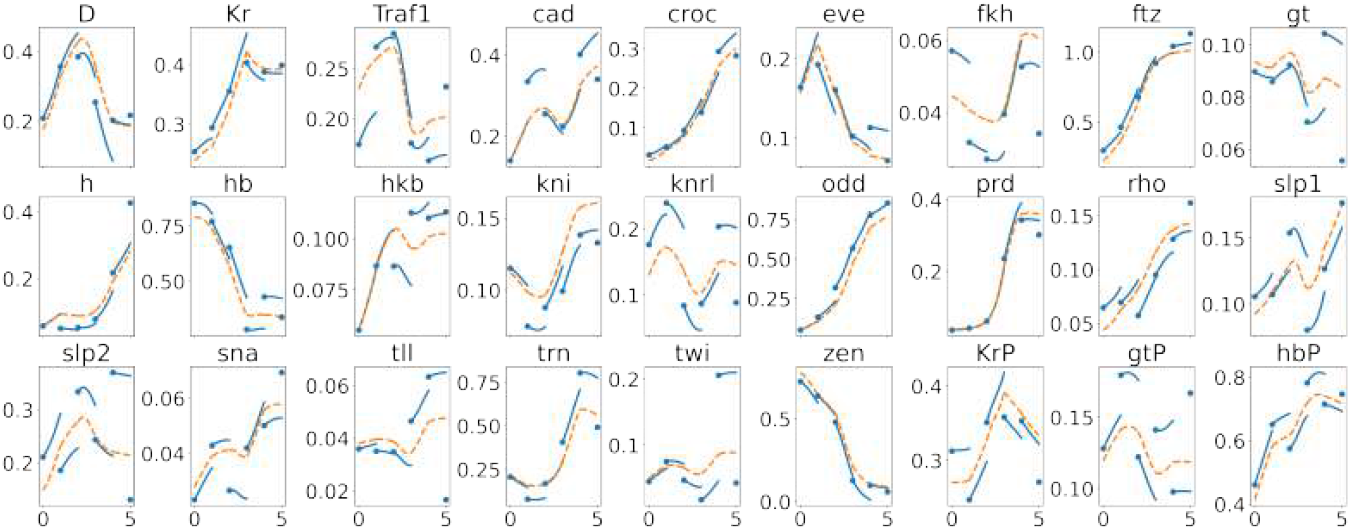
The trajectories produced by the trained neural network in the 27-dimensional gene space for one representative cell. It uses the VAE trained with 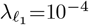 (see Table 1). Blue dots are observed expression at discrete times. Solid blue curves integrate in gene space starting from the observed data, eliminating reconstruction error at the segment start; dashed red curves decode the trivial latent dynamics by linear interpolation between points, incurring reconstruction error at both ends of the curves. Gene names are shown above each subplot; segments are not connected across boundaries.

In Fig 5, we compare mean absolute prediction errors across 27 fully measured genes between our black-box model, the Hill model, and the erf-weighted LAD models of (12). While the LAD quadratic model achieves the lowest raw error, our black-box approach remains competitive, and the Hill fits guided by the black-box dynamics offer a complementary advantage: they embed regulatory interactions directly in mechanistic Hill-type nonlinearities, rather than being limited to linear or quadratic couplings. Although the mean errors of our models are modestly higher, they produce interpretable functional forms that can be directly manipulated and biologically motivated, thereby providing a framework well-suited for hypothesis generation and experimental design.

**Fig 5.**
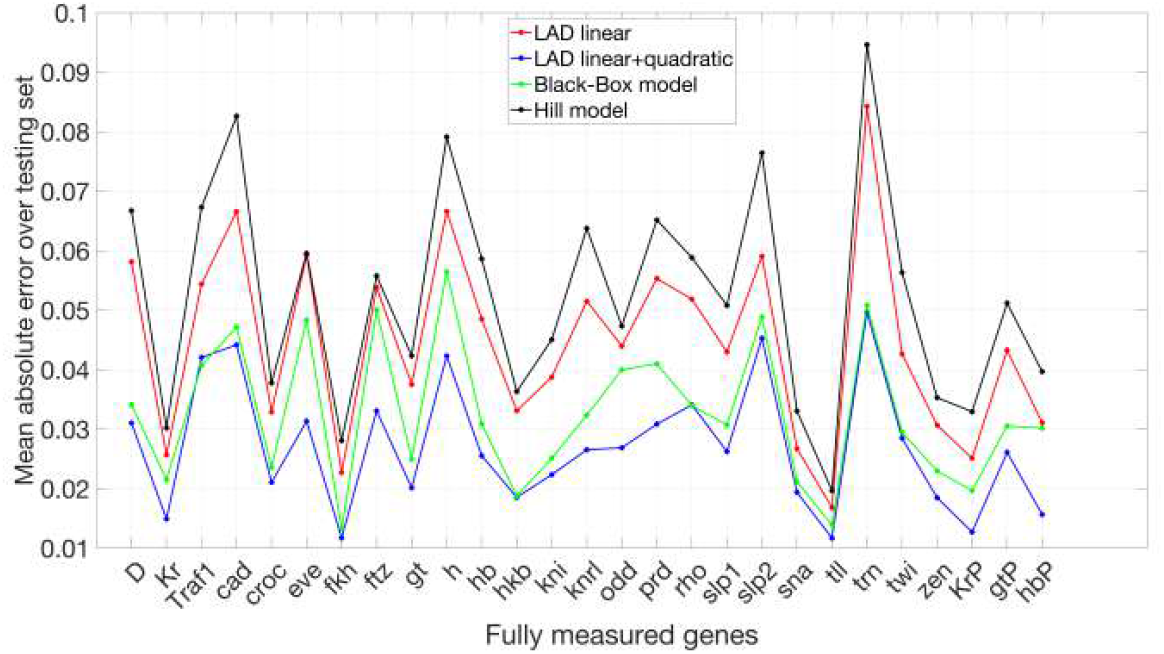
Mean absolute prediction error on the testing set. Comparison of mean absolute errors for our black-box model and the Hill model, alongside results reported in (12). The connecting lines are included solely for visual guidance and do not carry interpretive meaning.

Finally, we return to the full 99-dimensional gene space, train the black-box teacher, and fit the Hill model there, using VAE imputation to fill missing entries. In some settings the *teacher guided* Hill fit may underperform the *data-only* version; however, the deep-learning stage remains essential because its imputation enables derivative estimates and one-step targets where observations are absent, making the full-network fit feasible. Thus, the black box contributes to the mechanistic fit either by providing a useful teaching signal or, at minimum, by supplying the missing values needed to construct targets across all 99 genes.

To visualize how teacher guidance shapes downstream mechanistic fits, Fig 6 overlays continuous trajectories produced by the black-box model (VAE with latent neural ODE; blue) and the corresponding Hill-field rollouts (red) directly in the dimensional gene space between observed time points. Solid blue dots indicate measured expressions and hollow circles indicate VAE-imputed entries at the same states. Subplots titled in green are the 27 fully observed genes.

**Fig 6.**
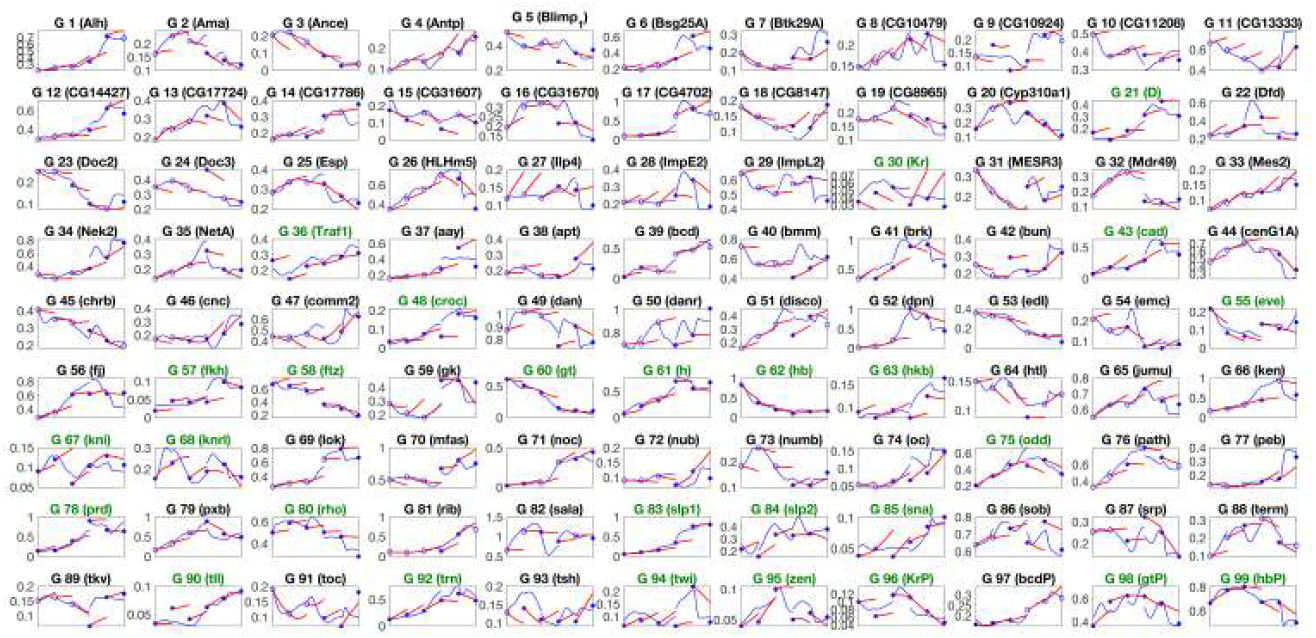
Black-box versus Hill-field trajectories in 99-dimensional gene space. Solid blue dots mark observed gene-expression measurements; hollow blue circles indicate values imputed by the VAE. Blue curves are the decoder applied to the latent neural ODE trajectories. Red curves are integrations of the corresponding Hill model trained from this black-box teacher. Subplot titles shown in green denote the 27 genes that are fully observed across all six time points.

### Universality of the Hill Model Inferred from Latent-Derived Dynamics

To assess the spatial universality of the mechanistic model inferred from our learned latent dynamics, we examined how restricting supervision to contiguous regions of the embryo affects prediction accuracy across the full embryo. This parallels the regional training experiments in (12), but here we apply it to Hill fields fitted to gene-space velocities produced by the black-box (hereafter abbreviated BB) teacher. For consistency, we did not retrain the VAE with a neural ODE but instead used the trivial latent linear dynamics together with the VAE pushforward to obtain gene-space derivatives.

We partitioned the embryo along either the anterior–posterior axis into three equal-sized regions (anterior, middle, posterior; AMP) or along the dorsal–ventral axis into dorsal, medial, and ventral (DMV) subsets (Fig 7). Each region covers roughly one third of the positional bins after test-set removal, i.e. about 33% of the training pool (the 90% of bins not in the fixed 10% test set). For each region, we trained a Hill field using only that region’s bins across all time points, following the same *ℓ*_1_-regularized training and relative-threshold pruning procedure used elsewhere. To ensure that differences across regions reflect supervision alone rather than variability in upstream representation learning, we did not retrain a separate latent neural ODE for each case. Instead, we derived gene-space derivatives directly from a fixed VAE pushforward with trivial latent dynamics, so that all regional Hill models are trained from a common, consistent source of velocities. For comparability, we also omit the final calibration step here, reporting raw regional fits without further adjustment.

**Fig 7.**
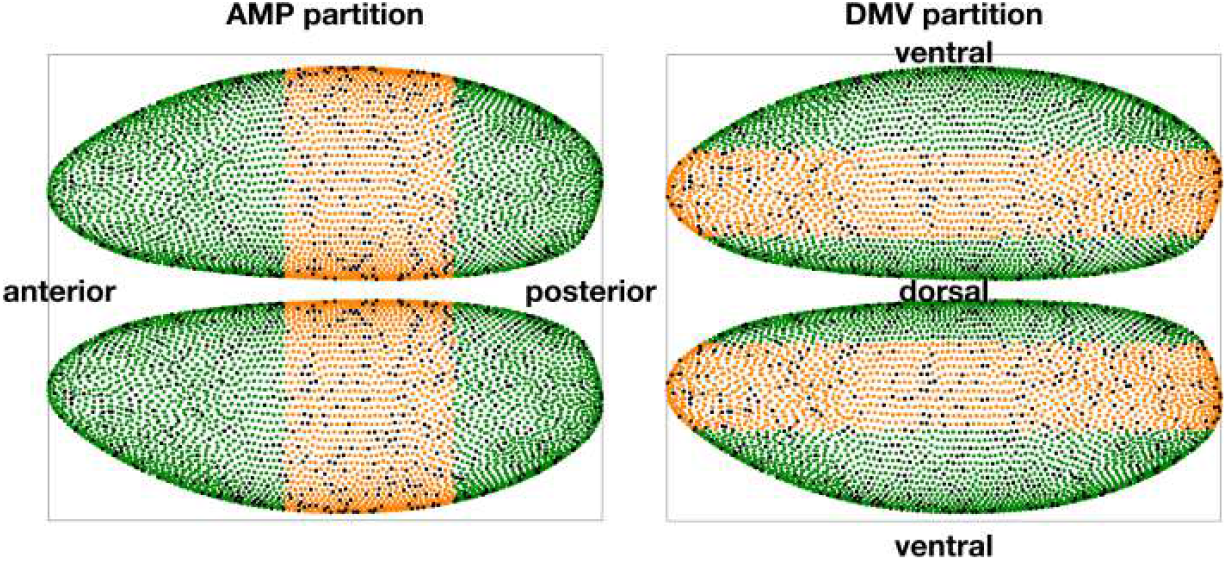
Spatial partitioning of the embryo into anterior–middle–posterior (AMP) and dorsal–medial–ventral (DMV) subsets. Cells are colored according to their regional assignment: anterior/posterior or dorsal/ventral in deep green, medial in orange. Black dots indicate the fixed 10% test cells that were excluded from training. These partitions define the subsets of cells used for region-specific Hill-model fitting. As a baseline, we also trained models on a random selection of 30% of bins uniformly distributed across the embryo (random-30%), matching the number of bins used in each regional subset.

As a baseline, we also trained “random-30%” models by uniformly sampling 30% of bins from the training pool—the same fraction as in each regional case (AMP or DMV). This ensures that both region-specific and random baselines use equal training size. To reduce variance from sampling, we repeated the random-30% selection with 10 different seeds and averaged the resulting errors. Prediction quality throughout this section is reported in terms of the relative error, defined as |*g −ĝ* |*/*(|*g* + *ĝ* |+ *ε*), where *g* denotes the ground-truth gene expression and *ĝ* the corresponding prediction obtained by integrating the Hill model. This normalization prevents highly expressed genes from dominating the error metric and ensures comparability across genes of different magnitudes.

Fig 8 and Fig 9 summarize per-gene prediction errors under the two regularization regimes. At *λ* = 10^*−*4^ (stronger regularization), all regional models achieve broadly similar accuracy, but the random-30% baseline provides a small yet consistent advantage. At *λ* = 10^*−*5^ (weaker regularization), this advantage becomes more pronounced: random-30% models outperform any single-region training, showing that broad coverage is particularly valuable when the network is less constrained and retains more couplings. These results indicate that balanced sampling across the embryo improves generalization, with the benefit becoming clearer as regularization is relaxed.

**Fig 8.**
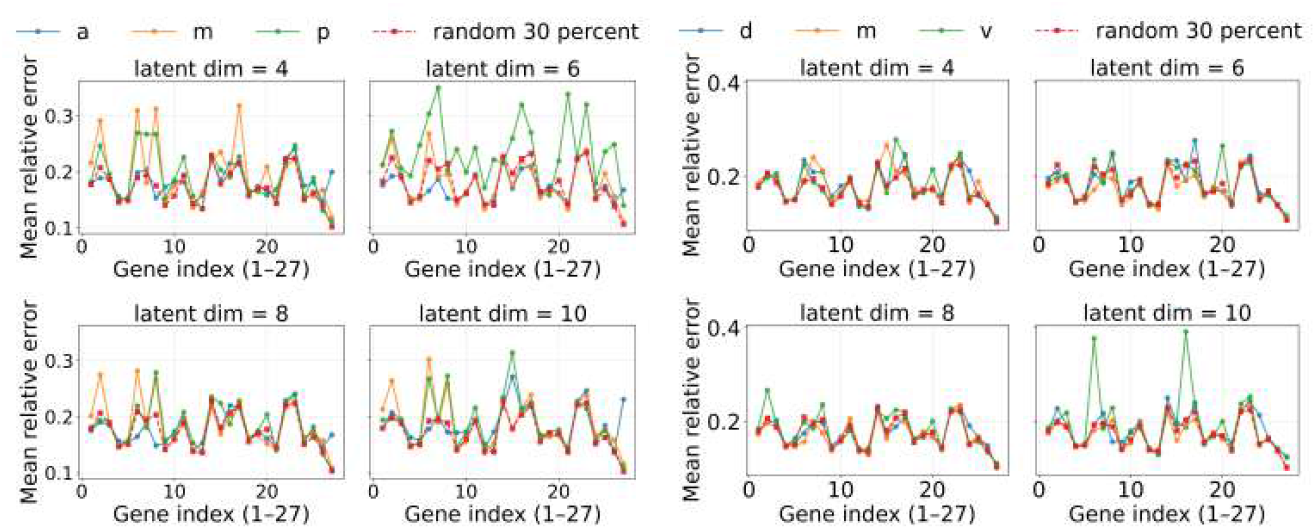
Universality of Hill-model fitting across regions for *λ* = 10^*−*4^ (stronger regularization). Mean relative error per gene for models trained on subsets of cells from either the anterior–middle–posterior (AMP, left) or dorsal–medial–ventral (DMV, right) partition, compared to models trained on randomly selected 30% of bins across the embryo (random-30%). Under stronger sparsity, the random-30% baseline already provides a modest advantage, suggesting that balanced spatial coverage improves generalization even when couplings are heavily pruned.

**Fig 9.**
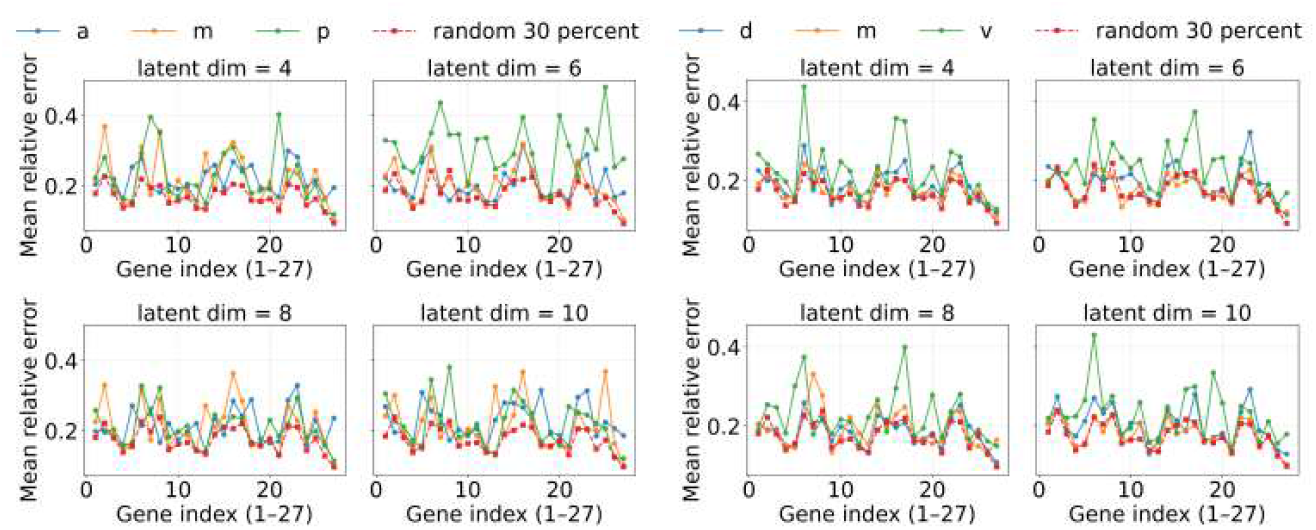
Universality of Hill-model fitting across regions for *λ* = 10^*−*5^ (weaker regularization). Mean relative error per gene for models trained on subsets of cells from either the AMP (left) or DMV (right) partition, compared to random-30%. With weaker sparsity, the advantage of random-30% becomes more pronounced: balanced coverage consistently outperforms any single-region training, indicating that broader sampling is especially valuable when the network retains more couplings.

As shown in Fig 10, the spatial overlays make clear that region-restricted supervision produces non-uniform error distributions across the embryo. For AMP training, anterior- and posterior-trained models tend to overfit locally, leaving elevated error bands in the opposite poles, while the middle-trained model shows weaker generalization in both anterior and posterior extremes (panels A–C, left). Similarly, under DMV supervision, dorsal- and ventral-trained models concentrate error in the opposite domain, and medial supervision underfits in both dorsal and ventral territories (panels a–c, right). By contrast, the random-30% baseline yields a more spatially uniform and consistently lower error profile across the entire embryo (panel D), suppressing the high-error pockets seen in the regional cases. This advantage is particularly evident in posterior and ventral domains, which are most prone to overfitting under single-region training. Crucially, the improvement is not restricted to a handful of genes or cells but extends across the full 27-gene set and all spatial positions, underscoring the benefit of balanced sampling for generalization.

**Fig 10.**
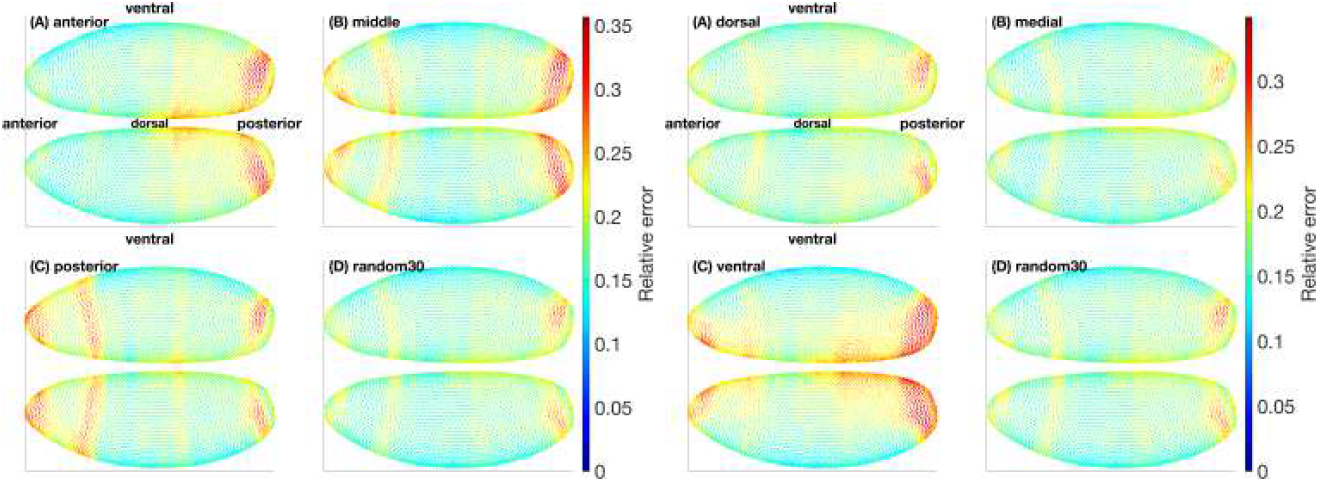
Spatial distribution of Hill-model errors for different regional partitions. Relative errors for the 27 fully measured genes are visualized on the embryo surface at latent dimension 10 with *λ* = 10^*−*4^. Panels (A–C) show models trained on individual regions (anterior, middle, posterior for AMP; dorsal, medial, ventral for DMV). Panel (D) shows the random-30% baseline. In both partitions, random-30% yields a more spatially uniform and consistently lower error distribution, especially in posterior and ventral domains where regional training tends to overfit.

### Mechanistic perturbations around *eve* via *gt* reveal interpretable, combinable controls

Having established the fitted Hill network, we next ask how its structure can be used to generate interpretable perturbation predictions. The sparse interaction matrix *V* defines a directed regulatory graph, visualized in Fig 11. Each node represents one of the 99 genes, with outline color indicating the sign of self-regulation, and edges correspond to nonzero couplings that survived pruning. Edges are drawn clockwise from regulator to target, with green indicating activation (*V*_*ij*_ > 0), red indicating repression (*V*_*ij*_ < 0), and transparency scaling with coupling strength. This graph-level view bridges the inferred dynamics with classical developmental biology by making explicit which genes act as activators or repressors of others. Among the resulting couplings, one particularly clear and biologically well supported interaction is the repressive influence of *giant* (*gt*) on *even-skipped* (*eve*). Because *eve* serves as a sensitive readout of upstream gap-gene inputs, this relationship provides a natural test case to illustrate how targeted perturbations—whether to state variables (e.g., gene knockout) or to specific coupling coefficients in *V* —reshape integrated expression trajectories and yield distinct transcript patterns.

**Fig 11.**
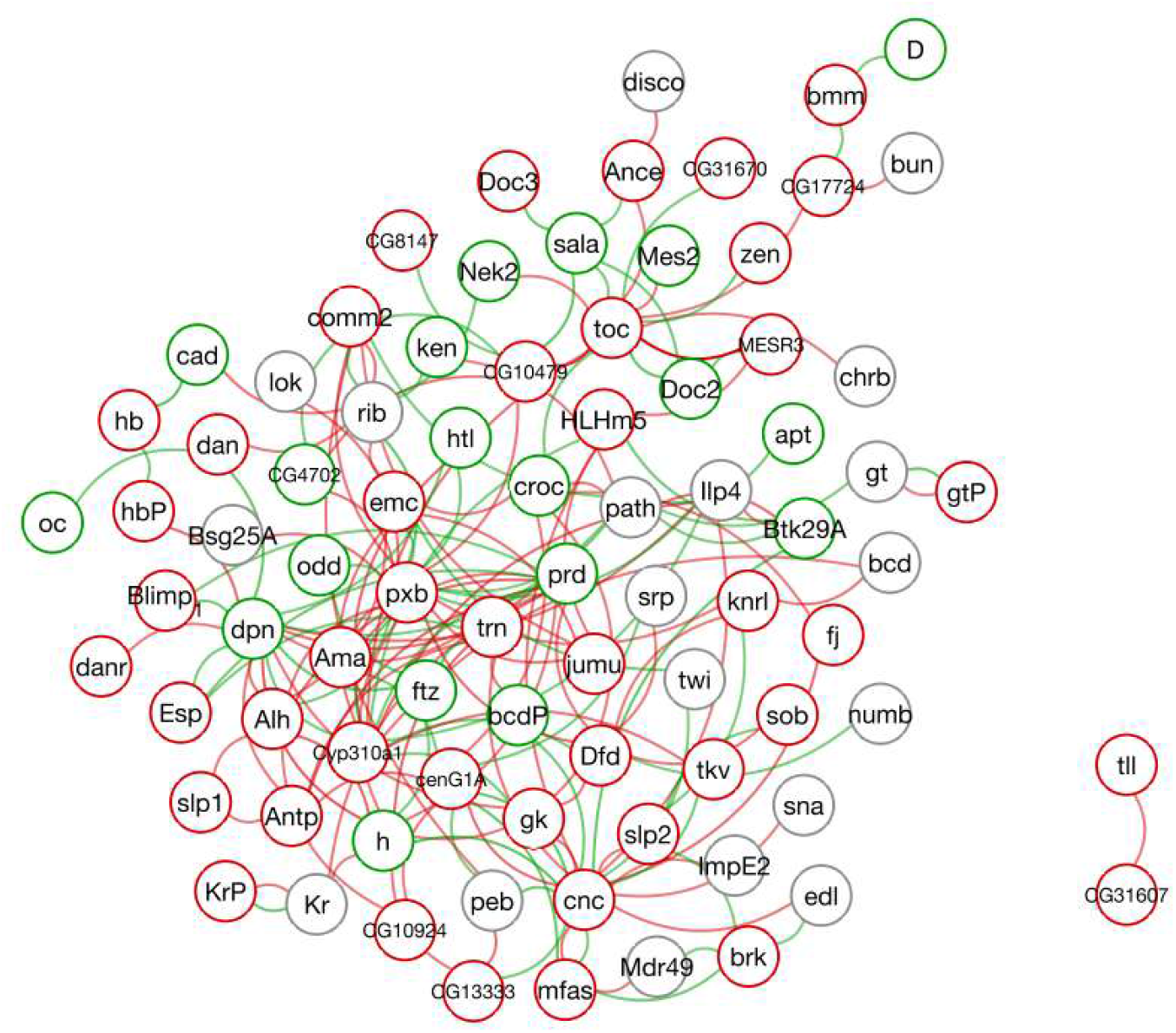
Inferred gene–regulatory network from Hill model couplings. Nodes denote genes, with outline color indicating the sign of self-regulation (green: positive, red: negative, gray: none). Directed edges represent fitted couplings *V*_*ij*_, drawn clockwise from regulator *j* to target *i*. Green edges indicate activation (*V*_*ij*_ > 0), red edges indicate repression (*V*_*ij*_ < 0), and transparency scales with coupling strength. This global network view links the Hill-field parameters to an explicit regulatory graph and motivates closer examination of specific interactions, such as the repressive effect of *gt* on *eve*.

We consider *even-skipped* (*eve*) as the readout and *giant* (*gt*) as a key upstream regulator, because their interaction is cleanly visible in our *t*=4 stripe pattern. During the blastoderm stage, *gt* acts predominantly as a repressor of *eve*, shaping the valleys between *eve* stripes. Biologically, *eve* is a pair-rule gene whose seven transverse stripes provide a sensitive readout of upstream gap-gene inputs, while *gt* is a gap gene with well-defined anterior–posterior domains that impose spatial repression on pair-rule enhancers. By *t*=4, which is about 10 minutes before gastrulation, the *eve* stripes are sharply refined and display approximate bilateral symmetry. In this regime, reducing *gt* input is expected to elevate *eve* primarily in inter-stripe regions while leaving stripe peaks comparatively less affected (17). Importantly, both *gt* and *eve* are fully measured across all six time points in our atlas, so the analyses here do not rely on imputation for these genes; moreover, the principal upstream regulators that influence *eve* are also largely in the fully observed subset, minimizing any dependence on imputed values for the relevant regulatory context.

To visualize how targeted edits propagate across the embryo, we show the *eve* level at *t*=4 across the full embryo in Fig 12. Colors encode *eve* abundance on a single, shared scale across all six subplots. Fig 12A presents the ground-truth atlas at *t*=4. Fig 12B to 12F display predictions of *eve* at *t*=4 obtained by integrating from *t*=2 using ground-truth (with imputation) gene values at *t*=2 under different conditions: Fig 12B integrates the BB teacher via decoder pushforward (VAE + latent neural ODE pushed through the decoder to gene space); Fig 12C integrates the Hill field; Fig 12D integrates the BB teacher with a knockout of *gt* (set to zero throughout *t*=2 → 4); Fig 12E integrates the Hill field with a *gt* knockout; and Fig 12F integrates the Hill field after setting the single coupling *V*_gt→eve_=0 while leaving all other parameters unchanged. Together, Fig 12A to Fig 12C establish reference distributions under a common color scale, whereas Fig 12D to Fig 12F show three targeted interventions—gene knockout in the BB teacher, gene knockout in the Hill field, and a single-edge edit in the Hill field—that reveal how specific upstream manipulations reshape *eve*’s *t*=4 pattern in a way directly attributable to modeled regulation. We select the *t*=2→4 window because it captures the refinement and growth of *eve* stripes; intervening substantially earlier or later is less informative for this readout.

**Fig 12.**
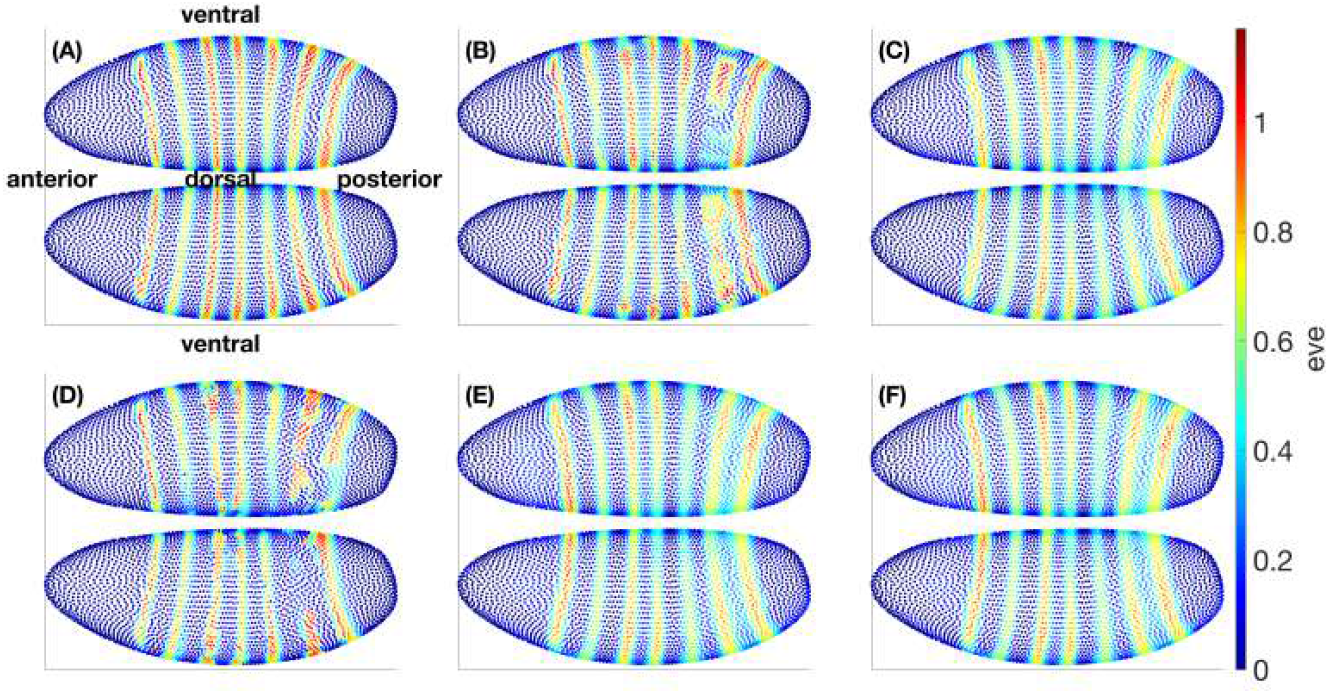
Absolute *eve* patterns under baseline and mechanistic/KO operations. (A) Ground truth at the target time. (B) BB baseline integration (decoder pushforward of the latent ODE). (C) Hill baseline integration. (D) BB with *gt* knockout. (E) Hill with *gt* knockout. (F) Hill with the single edge *V*_gt→eve_ removed while all other couplings are intact. A common color scale is used across all six panels to emphasize operation-driven differences.

To more clearly highlight the effects, Fig 13 shows the corresponding differences at *t*=4, computed as subtractions (D–A), (E–C), and (F–C) from Fig 12, where a single colorbar is used across all three panels. We plot black circles over the positional bins of the seven stripes (those with *eve* level above 0.3 at *t*=4). The dominant feature is a widespread positive shift (red) that weakens precisely at stripe crests and in their immediate inter-stripe gaps. This pattern is consistent with *gt* acting as a repressor of *eve*: removing *gt* or nullifying its specific influence broadly relieves repression, while regions already dominated by other inputs (stripe peaks) or locally balanced show weakened change.

**Fig 13.**
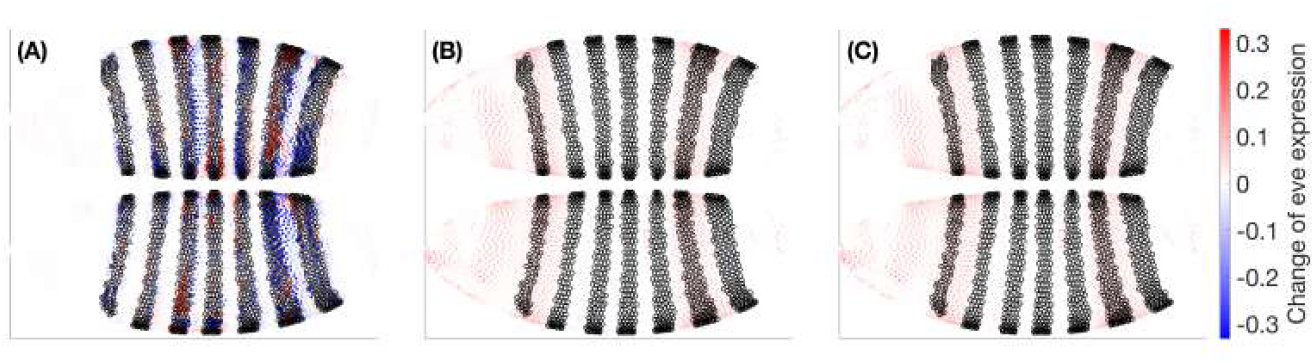
Operation–baseline subtractions reveal relief of *gt*-mediated repression. Signed differences (operation minus baseline) for: (A) BB *gt* knockout, (B) Hill *gt* knockout, (C) Hill removal of *V*_gt→eve_. All subplots share the same color scale. Broad positive changes away from stripe peaks indicate that reducing *gt* input generally increases *eve*, while muted changes at stripe maxima and specific gaps suggest compensating inputs and local balance.

A notable feature of Fig 13 is the close resemblance between Fig 13B (Hill-field knockout of *gt*) and Fig 13C (Hill-field edge edit with *V*_gt→eve_=0). The knockout sets *gt* to zero throughout the integration, removing both its direct influence on *eve* and any indirect, multi-hop routes (e.g., gt → *j* → eve). By contrast, the edge edit nullifies only the direct coupling *V*_gt→eve_ while leaving *gt* free to regulate other targets. The spatial concordance of the two Δ fields therefore indicates that, over the *t*=2 → 4 window, the dominant pathway by which *gt* modulates *eve* is the direct gt → eve link; putative indirect contributions via intermediates are comparatively small or cancel in aggregate.

This agreement strengthens the causal interpretation afforded by the mechanistic model: targeted edge-level edits isolate specific regulatory routes and, in this case, recapitulate the knockout’s effect on *eve* without altering the rest of the network. (This inference is conditional on the genes, time window, and operating regimes analyzed here; other contexts could reveal stronger multi-hop effects.)

Across models, the spatial signatures largely agree—supporting a shared causal interpretation—while differences are informative: the BB knockout Fig 13A tends to produce smoother, more diffuse changes, whereas the Hill-field contrasts Fig 13BC exhibit sharper, more localized structure that reflects explicit regulatory couplings. As expected, the edge edit Fig 13C mirrors the knockout’s sign pattern but with reduced amplitude, since it removes only the *gt* → *eve* interaction rather than the gene’s full influence. Because all panels share the same color scale, these amplitude relations are directly readable and highlight the practical advantage of the mechanistic model: it allows targeted, single-edge edits that isolate specific regulatory pathways—edits that are cumbersome to realize or interpret within a purely black-box dynamical model.

The patchwork of red and blue in Fig 13A (BB-knockout minus BB–baseline) reflects that a knockout in the BB teacher is not a clean removal of a single mechanistic influence but a global perturbation of the latent trajectory and its nonlinear readout. Concretely, setting *gt*=0 at *t*=2 changes the full 99-dimensional input that the VAE encoder maps into latent space, so the initial latent state—and thus the entire integrated path of the latent neural ODE—moves to a different region of the latent manifold where features entangle many genes. The decoder then converts that altered trajectory back to gene space through general nonlinear maps, with no sign constraint linking *gt* and *eve*; the local decoder Jacobian *J*_Dec_(*µ*) can point in different directions across positions, yielding increases in some locations and decreases in others. Because such a global zeroing pushes inputs away from the training distribution, the BB dynamics and decoder also extrapolate off-manifold, which can amplify non-monotonic, compensatory responses. Finally, the BB objective is pure predictive loss minimization, not pathway fidelity, so it can “balance” an imposed change through other latent directions to keep outputs close to typical values—again producing mixed signs. By contrast, Fig 13BC (Hill–knockout subtracts Hill–baseline and *V*_gt→eve_=0 subtracts Hill–baseline) edit an explicit term with fixed sign in the mechanistic ODE, so they predominantly show coherent increasing of *eve* where *gt* normally represses *eve*, with minimal sign flipping except near stripe peaks and narrow gaps where other regulators dominate.

## Conclusion

Under matched sparsity and calibration, Hill fields guided by BB-teacher, decoder–push-forward derivatives achieve equal or lower one-step forecast error than data-only (finite-difference) fits, with the largest gains at stronger *ℓ*_1_ regularization (Table 1). In global evaluation on the fixed 10% test set (608 bins), training on spatially restricted AMP or DMV regions underperforms training on randomly distributed subsets of comparable size, whereas random-30% supervision retains broad predictive accuracy (Figs. 8–9, 10). Mechanistic editing exposes causal pathways: for *gt* →*eve*, zeroing the single entry *V*_gt→eve_ closely reproduces a *gt* knockout over *t*=2 → 4, indicating that the direct link dominates multi-hop routes in this window (Figs. 12–13).

The same procedure used here to fit the Hill model also enables direct visualization of the inferred interaction structure as a gene–gene network (Fig 11). Such representations provide a bridge between computational inference and classical developmental biology by highlighting putative activators and repressors that can be compared with the literature and with future perturbation experiments.

Compared with BB knockouts—which perturb the latent state and propagate through the decoder Jacobian *J*_Dec_(*µ*)—the Hill field yields sharper, signed, and composable responses that are straightforward to interpret. Together, these findings support a division of labor: latent dynamics denoise and interpolate to provide smooth, decoder-consistent gene-space velocities, while the Hill field converts them into an editable, stable ODE that enables systematic, combinatorial perturbation studies.

## Discussion

This work demonstrates that learned latent dynamics and explicit mechanistic structure can be combined to obtain models that are simultaneously accurate, stable to integrate, and easy to interrogate. Using a VAE and latent neural ODE to interpolate sparse temporal observations and to supply smooth gene-space velocities via decoder pushforward simplifies subsequent identification of a sparse Hill field. Conversely, the mechanistic model exposes a signed interaction matrix *V* that can be edited directly, enabling controlled, pathway-level interventions that would be cumbersome to realize or interpret within a purely black-box system. The empirical comparison under matched regularization and calibration (Table 1) indicates that the latent stage acts as an effective variance-reduction device for velocities, particularly when the interaction budget is tight.

Our analysis of spatial supervision clarifies a complementary point: continuous latent dynamics do not replace the need for spatially diverse training bins. Under global evaluation on the fixed 10% test set, region-restricted training (for either the BB teacher–driven fit or the Hill student) degrades performance relative to randomly distributed subsets of comparable size. The random-30% baseline maintained accuracy close to that of the global model, and its advantage over region-specific supervision grew stronger under weaker sparsity constraints (Figs. 8–9). In practice, embryo-wide regulatory structure requires embryo-wide coverage; gains from representation learning are maximized when spatial sampling spans the regimes in which predictions will be made.

Mechanistic editing around *gt*→*eve* illustrates how explicit couplings turn qualitative intuition into quantitative, testable hypotheses. Nullifying a single edge in the Hill field closely reproduces the effect of a *gt* knockout over *t*=2→ 4, suggesting that the direct pathway dominates multi-hop routes in this regime (Figs. 12–13). Because edits are applied to explicit entries of *V*, multiple interventions compose without retraining, converting an exponential black-box “what-if” space into a tractable sequence of controlled ODE integrations.

Several design choices improved learnability and stability. Including spatial coordinates in the encoder but not in the decoder target preserved spatial context without forcing the decoder to reproduce geometry. Decoder-smoothness and directional-alignment terms traded a small amount of raw reconstruction accuracy for better-conditioned *J*_Dec_(*µ*) and more coherent gene-space flows. Scheduling the Hill loss from derivative matching toward one-step prediction reduced overfitting to local slopes while ensuring consistency under integration.

Several limitations need to be considered. The atlas has only six time points, with many unmeasured genes, so even mask-respecting imputation can bias the teaching signal; this temporal sparsity and partial observability reduce identifiability, as multiple Hill parameterizations can explain similar flows even after *ℓ*_1_ pruning and calibration, and some inferred couplings may reflect effective rather than direct regulation. Pushing dynamics through the decoder depends on accurate local Jacobians; if the manifold is curved, the first-order pushforward may distort higher-order effects. The present functional form uses single-input Hill terms, which cannot capture combinatorial logic such as requiring both regulators (AND) or either regulator (OR); parameters are shared across all 6,078 bins, though real thresholds vary along anterior–posterior and dorsal–ventral axes. Performance is reported mainly on reconstruction and one-step prediction error; it would be informative to also analyze stability near fixed points, detect limit cycles, assess sensitivity to perturbations, and perform temporal hold-outs. Finally, we have not yet provided matched benchmarks against dynamic methods such as SCODE, GRISLI, GpFuse, or vector-field estimators, and the black-box ODE is not explicitly constrained to preserve non-negativity or boundedness.

Our future plan focuses on addressing these limitations without altering the core framework. To make uncertainty explicit, we could adopt Bayesian sparsity for the interaction matrix and report credible intervals for edges and trajectories, while propagating encoder, decoder, and latent-ODE variance into the student fit. We could also empirically validate the pushforward linearization by comparing *J*_Dec_(*µ*)*f*_*θ*_(*µ*) to finite-difference slopes computed along decoded micro-rollouts over short latent arcs; agreement scores would then gate which signal (pushforward versus decoded slope) supervises each sample. Higher-order synergy could be approximated by additional hidden nodes or by generalized Hill functions with multi-input denominators. A simple extension would let selected parameters depend linearly on spatial covariates or include region-specific random effects. Evaluation could be broadened with temporal cross-validation and lightweight dynamical diagnostics (local stability spectra, sensitivity to parameter perturbations, simple limit-cycle probes). Physical plausibility can be strengthened by positivity-preserving or projected integrators for the black-box ODE to enforce non-negative production and practical boundedness.

Beyond these steps, we plan to adapt the likelihood to single-cell counts by replacing the Gaussian reconstruction with negative binomial (NB) or zero-inflated negative binomial (ZINB) formulations to handle over-dispersion and dropout; introduce hierarchical priors on interaction weights to share information across genes with similar cis-motif enrichments; and perform posterior predictive checks by simulating the fitted mechanistic ODE beyond the observed window and comparing emergent spatial stripes with classical patterns. As a complementary extension, we could move from ODEs to spatially explicit reaction–diffusion partial differential equations (PDEs) on the embryo surface, pairing a neural PDE teacher with a sparse mechanistic reaction–diffusion student to capture diffusion/advection and boundary conditions while preserving editability.

In summary, the teacher–student design delivers variance-reduced, decoder-consistent derivatives together with a sparse, editable Hill field that supports signed, composable perturbations—turning black-box accuracy into causal, designable models. The incremental extensions outlined above provide a clear, testable path to strengthen the framework—adding quantified uncertainty, linearization checks, modest combinatorial and spatial expressivity, richer evaluation, and count-aware adaptations—while preserving its core practicality and interpretability. As datasets expand in scale and modality, this combination of smooth latent dynamics with an explicitly editable mechanistic layer offers a broadly applicable template for extracting robust regulatory insight from high-dimensional single-cell atlases.

## Supporting information

**S1 File. Supplementary Information**. Detailed methods, mathematical derivations, hyperparameter settings, ablation studies, and additional tables supporting the main text.

## Acknowledgments

This research was supported by the Intramural Research Program of the National Institute of Diabetes and Digestive and Kidney Diseases (NIDDK) within the National Institutes of Health (NIH). (Project 1 ZIA DK075146-06. V.P. received this grant.) The contributions of the NIH author(s) are considered Works of the United States Government. The findings and conclusions presented in this paper are those of the author(s) and do not necessarily reflect the views of the NIH or the U.S. Department of Health and Human Services.

